# Moose or spruce: A systems analysis model for managing conflicts between moose and forestry in Sweden

**DOI:** 10.1101/2020.08.11.241372

**Authors:** Oskar Franklin, Andrey Krasovskiy, Florian Kraxner, Anton Platov, Dmitry Schepaschenko, Sylvain Leduc, Brady Mattsson

**Affiliations:** IIASA, International Institute for Applied Systems Analysis, Ecosystem Services and Management Program, Austria; University of Natural Resources and Life Sciences, Austria

## Abstract

Sweden has the world’s highest density of moose (*Alces alces*). Moose is not only a valuable game species; it also causes forest damages and traffic accidents. To avoid moose browsing, foresters respond by planting spruce (*Picea abies*) to an extent that reshapes the forest landscape with impacts on both production and biodiversity. To address this problem and maintain a healthy moose population in balance with the other interests, an adaptive management based on the knowledge and experiences of local hunters and landowners is advocated. However, the different stakeholders do not agree on what is an appropriate moose population, which leads to conflicts that are hard to resolve. A key problem is that it is very difficult to encompass and foresee long-term consequences of different options for moose hunting and forest management. This makes it challenging to form coherent strategies that integrate different sectorial interests at a national level. To address this issue, we have developed a systems analysis framework for integrated modeling of the moose population, forestry, and their interactions and consequences for biodiversity. We analyze the short and long-term consequences for multiple scenarios of moose hunting and forest management. Based on the results we elucidate and quantify the trade-offs and possible synergies between moose hunting and forest production. This analysis can be used to support better informed and more constructive discussions among the stakeholders in the Swedish forest sectors, and to support policies for long term sustainable forest and moose management.

## Introduction

### Background and importance of moose management

Sweden has the world’s densest moose (*Alces alces*) population in relation to the forest area. During winter moose consume large amounts of Scots pine (Pinus sylvestris), which leads to local pine stands being severely damaged. The impact of moose browsing on the Swedish forests are substantial and according to the Swedish National Forest Inventory’s data from the year 2009-2013, approximately 42% of the scots pine in Sweden suffers from moose browsing damages (Skogsstyrelsen, 2016). Estimates of economic loss in forestry has proven difficult and studies therefore differ substantially in their results. For example, calculations made in 2004 showed that the browsing pressure of pine forests by moose, was expected to cause a 30-80 million SEK loss of income for forest owners in Sweden each year (Glöde et al. 2004). Calculations also showed that the browsing pressure would result in additional losses in quality of approximately 500 million up to 1,3 billion SEK in 30-50 years (Glöde et al. 2004). This should be compared to a report presented by the National forest agency (NFA) in 2007 concluding that the costs inflicted to forestry amounts to about 650 million SEK on an annual basis (Ingemarson et al. 2007). Due to the potential biological and economic consequences, the management of the moose population is evidently important.

### Current moose management and stakeholders

The moose population in Sweden has historically fluctuated considerably, from a situation near extinction to a state of overabundance. With the intent to solve some of the ecological and social problems present in the moose management, the Swedish government introduced a new moose management system in 2012 (SEPA 2018). Management is now supposed to be carried out in an adaptive, ecosystem- and locally based way, were the knowledge and experiences of hunters and landowners are used to manage the species. A new level of responsibility has been installed called moose management areas (MMA). Each MMA is generally intended to include one distinct moose population and to be ≥50 000 hectare in size. The County administrative board (CAB) is responsible for dividing their respective county into an appropriate number of MMA ‘s and as each area is supposed to be confined by natural barriers, the range of an MMA can extend beyond county borders.

The MMA’s are expected to facilitate the connection between the local hunters, landowners and the CAB. The MMA is governed by a moose management group (MMG). The MMGs most commonly consists of three hunters’ representatives and three landowners’ representatives, who are supposed to represent the interests of their respective organizations (Ingemarson et al. 2007). Exceptionally, in some parts of northern Sweden one of the hunters’ representatives is replaced by a Sami representative. The representatives are nominated by the hunters’ and landowners’ organizations and are then elected by the CAB. According to the proposition (2009/10:239), the MMG representatives are supposed to have, among other things, knowledge regarding forestry and hunting. They are supposed to represent the interests of all local hunters and landowners and are expected to formulate goals for aspects such as the winter population of moose, the amount of damages on forests and crops which are acceptable, harvest levels etc. Moreover, the MMGs are responsible for having consultations with the moose management units (MMUs) and developing a moose management plan (MMP) for the MMA which can be harmonized with the MMPs produced on an MMU level. Thus, responsiveness is an important part of the multileveled system. Furthermore, the Swedish government believe that, since the landowners are responsible for the care of the forests, they are to be given a stronger position in the MMG’s. Therefore, one of the landowner’s representatives is appointed chairman and obtains a casting vote. Consequently, should the voting end in a tie, the landowner’s chairman can use the casting vote and determine which decision should be made. However, a moose management plan must be approved by the CAB before it is finalized and available for use. Subsequently, the chairman does not have full authority to determine how the MMP is designed. The problem with the current moose management system is that both the landowners and the hunters have the same overall goal – a moose population in balance – however, they have not the same opinion on what that is.

### Goals and scope of the project

In order to alleviate the problem of diverging goals of the different stakeholders (hunters, landowners, county boards, SEPA, NFA), stakeholders need better tools to use as a base for more constructive discussions. In addition, effects on common or external aspects, such as biodiversity and traffic accidents, must be considered to avoid tragedy of the commons effects.

The goals of this project were to investigate the possibilities to develop modeling tools that can help improve Swedish moose management. This includes the following key tasks:

1. Identifying the current management problems and their causes
2. Clarifying and prioritizing the modeling needs and requirements
3. Acquiring the information and data necessary to construct the models
4. Developing prototype models for the key components of the system, forests and moose
5. Developing a prototype integrated model for the whole system for one MMA
6. Analyzing and evaluating the models’ capacity to address relevant questions for moose management.

The prototype integrated model should provide a capacity to understand the whole system consequences of different policy and management options for the interaction between forestry and wildlife (especially moose), other ecosystem services, biodiversity, and traffic accidents. It should be able to provide projections to improve and explain policy alternatives from the perspective of the different stakeholders i.e. SEPA, the Forest Agency (NFA), the county boards, and hunters and landowners in each MMA. In addition, the project is expected to provide knowledge and capacity in the field of wildlife management and modeling for all participants, improve methods and strategies for interactions between policy makers and researchers, and strengthen the collaboration between Sweden and IIASA.

Whereas the long-term goals of this work is to take account of a wide range of ecosystem and societal aspects of wild-life management for the whole of Sweden, this pilot project serves to demonstrate the feasibility of the approach by focusing on the limited but important interactions and trade-offs between forest- and moose management in one MMA. The model will produce spatially explicit estimates of forests and forest production, moose population, and felled moose for relevant scenarios for forest management and hunting. The results will be evaluated to quantify and illustrate potentials and tradeoffs between different objectives.

## Methods and data

### Overall framework

As illustrated in Fig. 1, there are dynamic, two-way interactions (feedbacks) between the moose population and the forests. In simple words, the moose browse forests, which affects forest growth, which feeds back on moose food availability and moose population dynamics. This puts some requirements on the modeling: The forest and moose models must be integrated. In addition to the output relevant for forestry, the forest model must produce the output relevant for the moose: species, height of trees, edible biomass, and effects of browsing on subsequent tree growth and wood volume production. The forest model must also produce relevant output for modeling biodiversity, such as dead trees. In addition to the dynamics of the moose population, the moose model must predict the effects on the forests, i.e. fraction of trees browsed per area.

**Figure 1.**
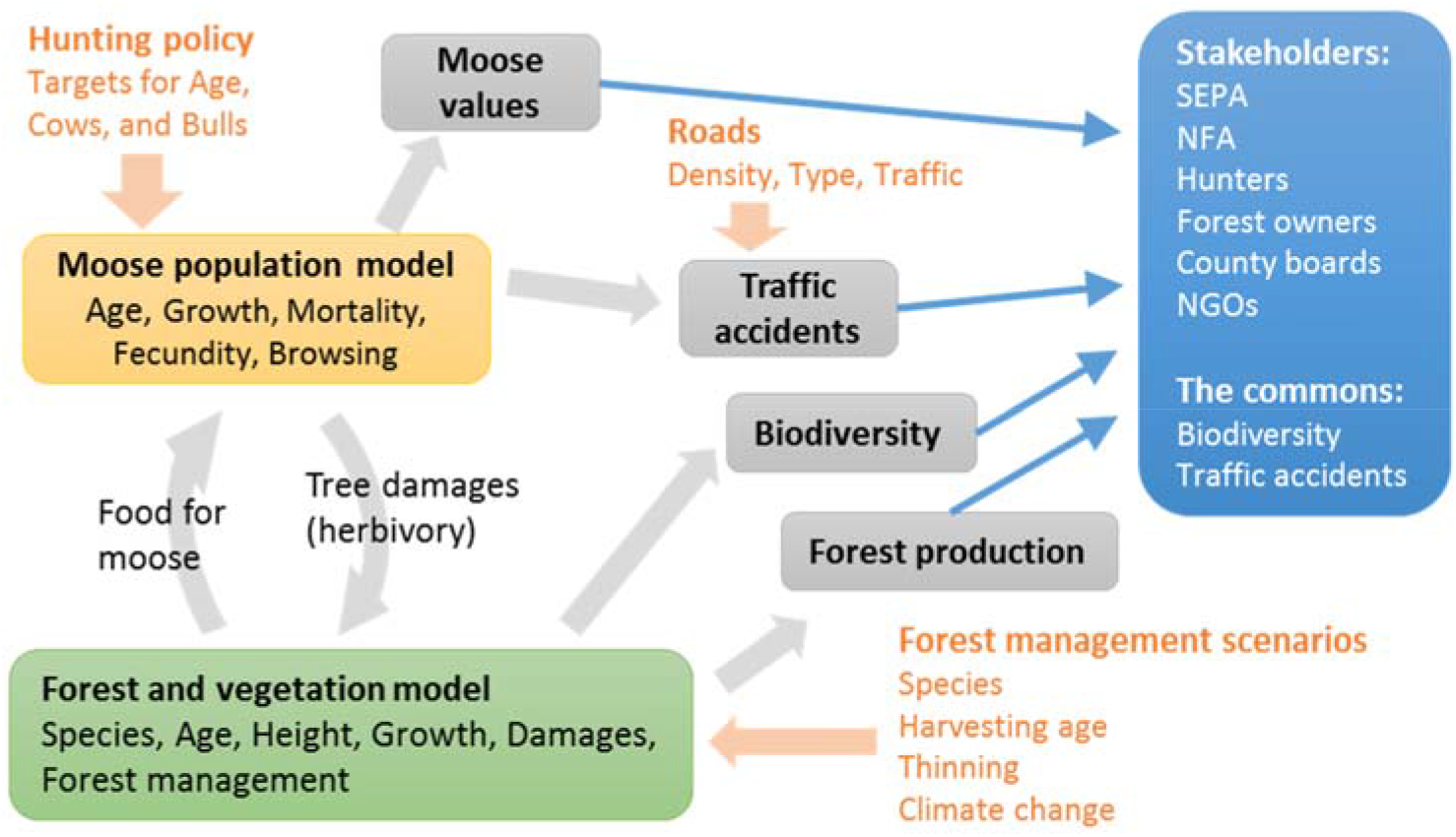
Overview of the main components of the analysis. The orange text shows the input variables that can be controlled by management or are externally supplied.

The appropriate spatial scale and resolution of the forest model depends on the resolution of forest spatial information and on computational limitations. To be able to flexibly adapt to future data availability and computational demands, the spatial model is designed to have adjustable resolution, i.e. size of the grid cells (Fig. 2). The spatial extent of moose populations in the model is initially defined as one population per MMA, but can be adjusted to any other area if needed.

**Figure 2.**
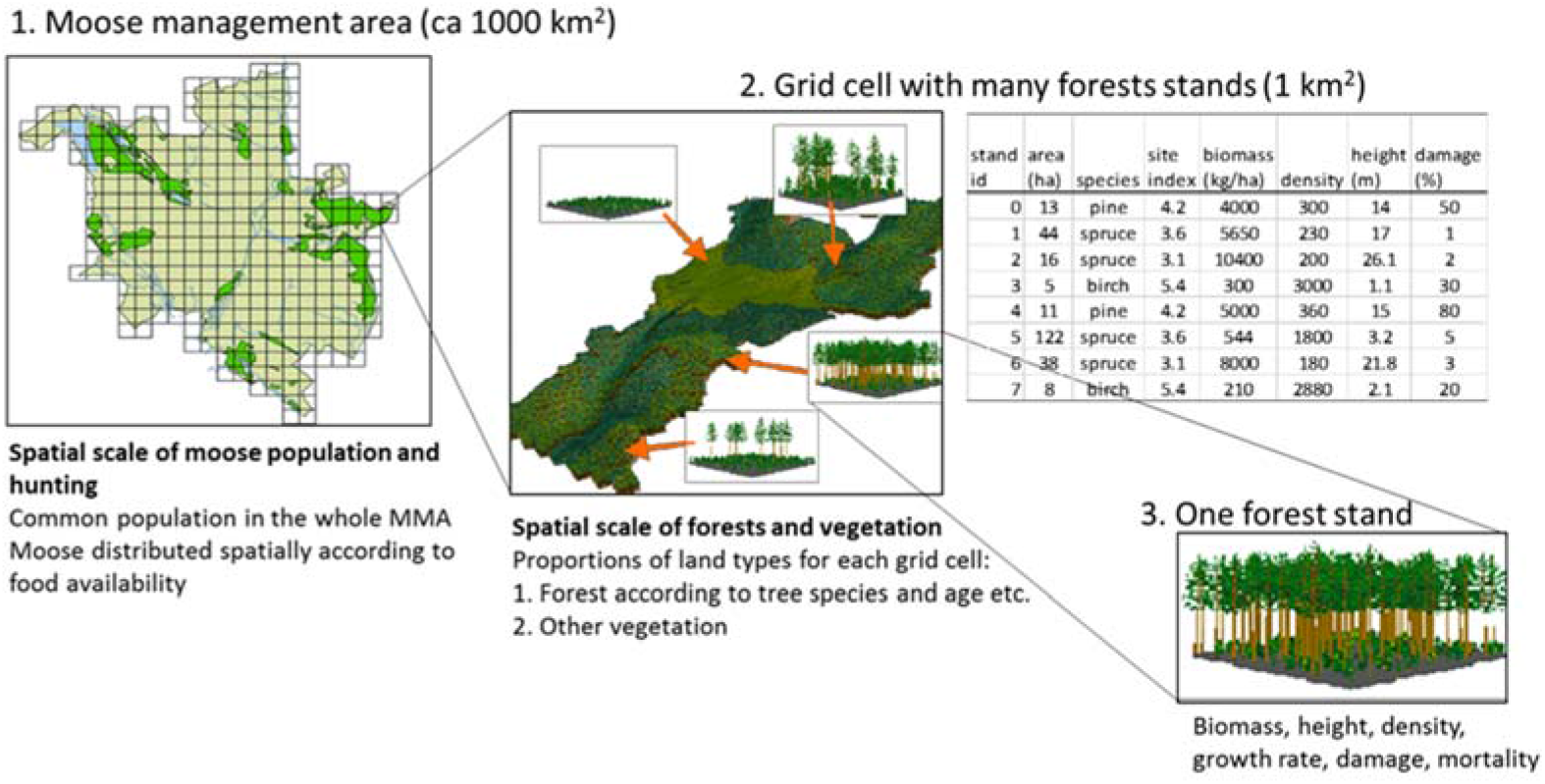
Spatial scales of the moose and forest sub models

### Forest model

#### Forest growth and management

To strike a good balance between computational speed and realism, we developed a stand-based forest model that describes the size and growth of biomass and height of the average tree, and the number of trees in each forest stand (Fig. 2). Biomass and height growth functions, as well as baseline mortality functions, for all relevant tree species in northern Eurasia has been derived based on extensive measurements (Shvidenko et al. 2008). Growth rate depends on site productivity (similar to the “site index” often used in forestry), which represents the integrated effect of all site factors (soil, water, climate) on growth. This site productivity parameter can also be used to model effects of climate change, rising CO_2_, and management such as fertilization.

To ensure that forest composition and growth is realistically modelled, observed data in each grid cell on species, age, biomass, and height (SLU 2018) is used to initialize the model and to estimate the site productivity based on the observed relationships between height and age in each cell (Fig. 3).

**Figure 3.**
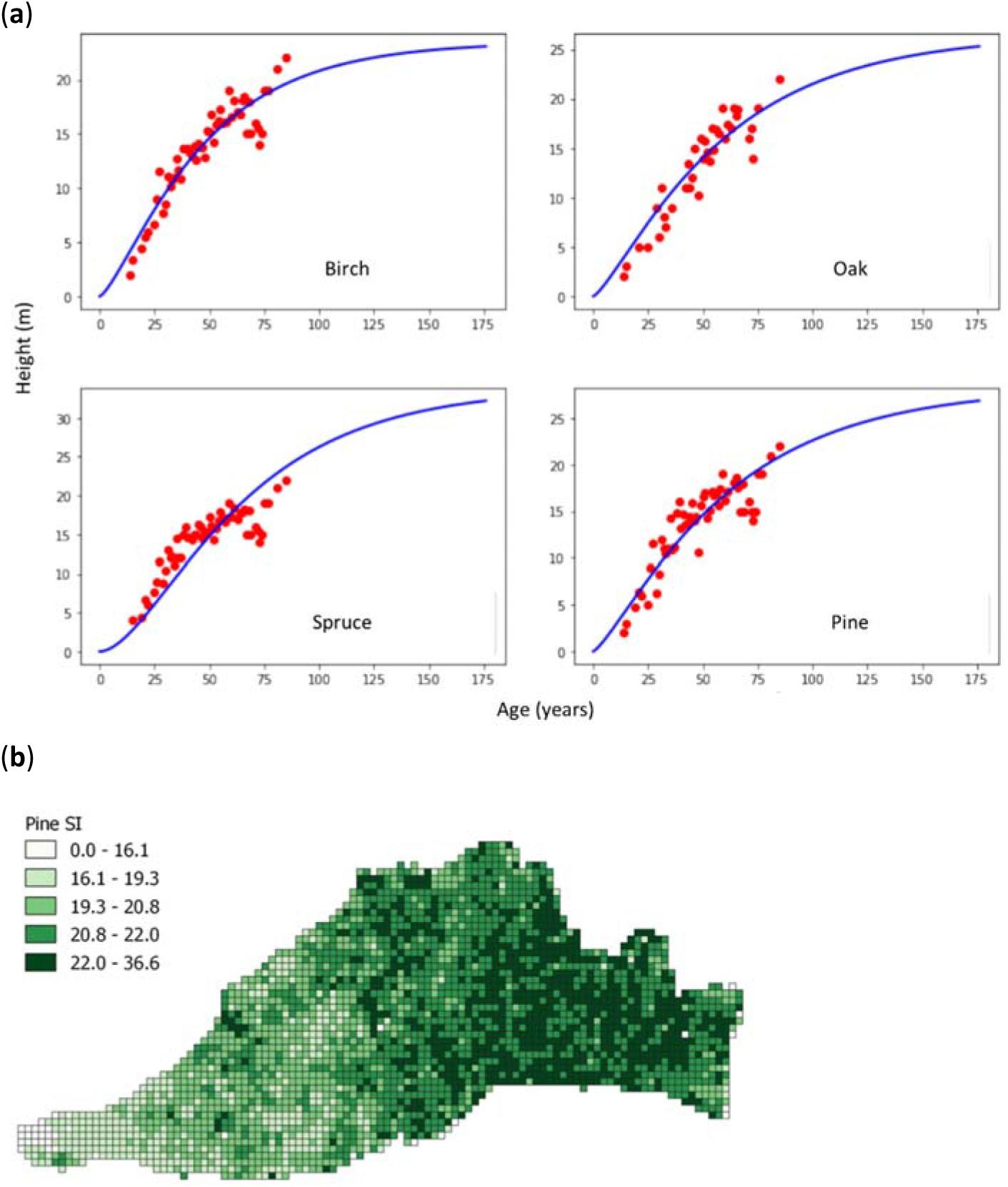
(**a**) Site productivity for a grid cell is estimated by fitting the forest growth model (blue line) to forest observations (red points). (**b**) Spatial distribution of estimated productivity in the moose management area studied (location shown in Fig. 6) measured as site index (SI, expected height at age = 100) for pine. Each cell is 1 × 1 km.

Stand density is estimated based on observed biomass compared to maximum biomass (maximum number of trees of a given size per area that can coexist) based on extensive historical observations for each species (Shvidenko et al. 2008). Thinning is modelled by reducing stand density which also increases growth and reduces mortality of the remaining trees (Franklin et al. 2009, Franklin et al. 2012). Harvesting and the tree age at harvest can be modelled based on different assumptions: empirical information on silvicultural practices in the area, estimation based on observed age distributions, optimization of biomass or volume production. Initially, we will estimate rotation time and thinning practice based on observed age distributions where possible, and otherwise apply fixed thinning and harvesting ages (70 years) for all stands in our test MMA.

#### Moose and forest interactions

The moose prefer certain tree species to others (broadleaved trees > pine >> spruce (*Picea abies*), (Shipley et al. 1998, Bergqvist et al. 2018)) and this is used to distribute the moose activity (browsing) in the landscape, according to an ideal distribution (Bergqvist et al. 2018). Moose eat leaf and twig biomass that they can reach, which is approximately up to a height of 3 m. This is used to calculate the food biomass available for moose browsing as a function of tree size (Kalén and Bergquist 2004). Moose browsing will be estimated based on the needs of the moose population (≈ 10 kg food per day and moose) unless it is limited by food availability.

Moose browsing reduce the biomass of trees and cause mortality. It also reduces the growth rate of the forest due to both damages on the individual trees and effects on size structure and species composition of the stand. This will be modelled by a damage index that reduces growth rate. The damages index increases with browsing and, in the absence of browsing, decreases over time as trees recover. Parameters for these functions were estimated based on long term moose browsing and exclusion trials (Pettersson et al. 2010) (Fig. 4)

**Figure 4.**
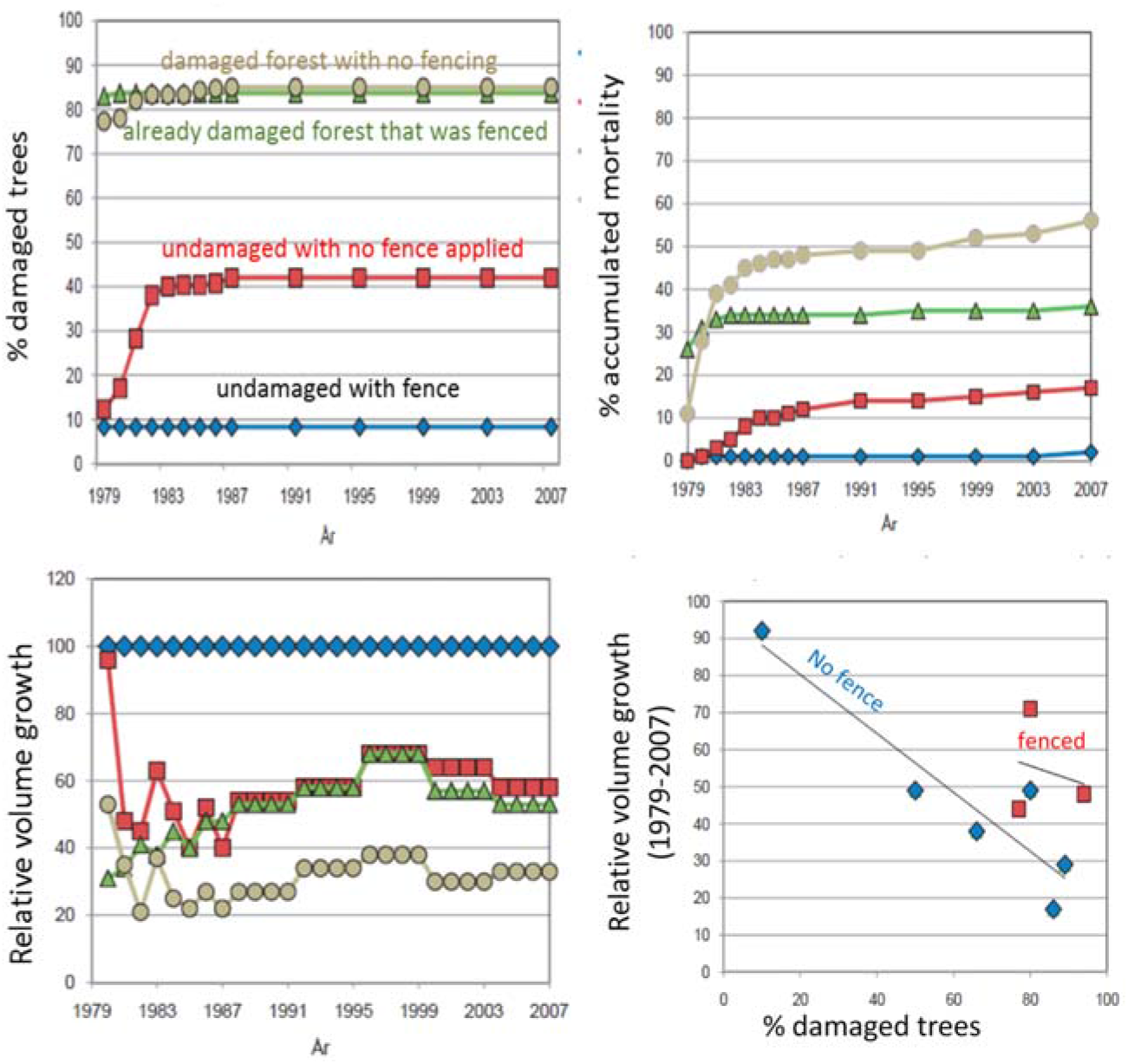
Example of effect of damage on stand volume growth and mortality from long term moose browsing and exclusion trials in pine forests. Re-drawn from (Pettersson et al. 2010).

### Moose model

We have developed an age- and sex-structured population model for moose, which includes growth, mortality and fecundity depending on age according to observations (Kalen 2018). Fecundity also depends on food availability and declines in proportion to food deficiency if food availability is less than the baseline food demand (Fig. 5a). Mortality includes death from natural causes (depending on age, Fig. 5b), traffic accidents, and hunting. Hunting is modelled in terms of the fraction of moose shot of each age class for bulls, cows and calves.

**Figure 5.**
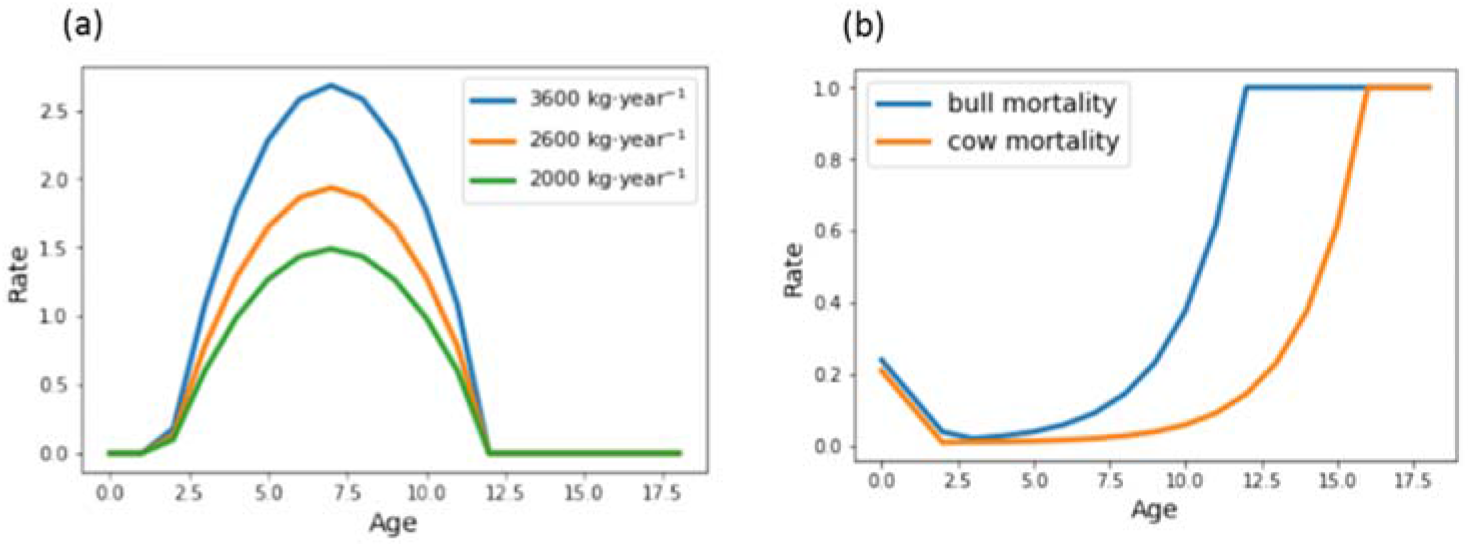
(**a**) Moose fecundity (maximal number of calves produced per cow per year) for maximum (3600 kg per year) and lower levels of food consumption. (**b**) Natural mortality (per year)

In this moose model, the moose population will grow until it is limited by hunting or by reduced fecundity due to food deficiency. In future versions, the individual level moose model will be developed to include food consumption, mortality and growth linked to moose weight, and also to account for climatic factors, such as the length of the winter.

### Model demonstration

To provide a proof of concept we aimed to evaluate the ability and usefulness of the integrated model to address questions relevant for moose management. This evaluation was done by running the model 20 years into the future for a number of moose and forest management scenarios and then evaluating the outcomes in terms of moose population, felled moose, and forest stocks, including emerging relationships and trade-offs between different goals (and respective stakeholders). Importantly, the purpose of this exercise was not to provide accurate and specific predictions to be used in practice but merely to test the model’s capacity to describe the system, address the relevant questions, and produce the relevant outputs. Thus, although we aimed for realistic model parameterization and scenarios, the model has not yet been validated and its output should be interpreted with care. We expect to produce reliable results for practical applications only after comprehensive testing and validation of the model in the next phase of the project.

### Scenarios

To test the model, we modelled different hunting and forest management scenarios (Table 1) in the moose management area (Älgförvaltningsområde) 8 in Västra Götalands län (Fig. 6). In all scenarios we simulate a time period of 20 years.

**Table 1.** Scenarios

| Forest management scenarios | Moose management scenarios* |  |
| --- | --- | --- |
| **BAU:** No species changes. | **Target population scenarios** |  |
| **Pine preference (Pine+):** Pine replanted + additional planting of pine in 5% of the previously non-pine areas. | Moose/km<sup>2</sup> | Moose in MMA |
|  | 0.63 | 1500 |
|  | 0.84 | 2000 |
|  | 1.05 | 2500 |
| **Spruce preference (Spruce+):** Spruce preference + additional planting spruce in 5% on the previously non-spruce harvested areas. | 1.27 | 3000 |
|  | 1.69 | 4000 |
|  | **Low hunting scenario** (10% per year) |  |
|  | **No hunting scenario** |  |

\*In all scenarios 50% of felled animals are calves and hunting is adjusted to keep the population at 65% cows

**Figure 6.**
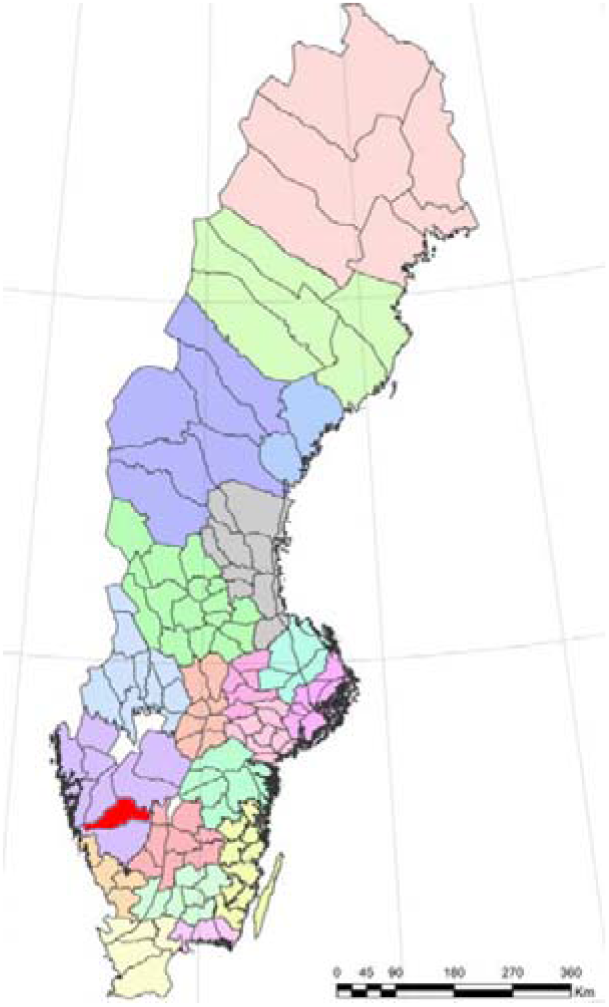
Map of the moose management areas (MMAs, n = 149) in Sweden. MMA boundaries are delineated by the black lines. Colours indicate county administrative boards (Län). The modeled MMA is Älgförvaltningsområde 8 in Västra götalands län, marked in red and has an area of 237000 ha.

For forest management we applied three different planting scenarios. In the business as usual scenario (BAU) the model always re-plant each area after harvest with the same species that was harvested. In the pine preference scenario (Pine+) we always re-plant harvested pine but also plant pine on 5% of harvested previously non-pine areas. In the spruce scenarios (Spruce+) we do the same as in the Pine+ scenario but for spruce (Table 1). Harvesting age and thinning practices were estimated based on the observed age distribution of the forests in the MMA. For example, a steep decline in the frequency of stands around 90 years age, indicate that this is a common harvesting age, and a steep decline of in volume per ha of around 20% at around age 45 indicates 20% thinning at age 45.

Moose scenarios are defined by a given target population in the MMA, which is maintained by applying hunting adaptively, i.e. if the population is higher than the target, hunting is increased and vice versa if the population is lower. We also modelled a low-hunting and a no-hunting scenario (Table 1). In order to maintain a productive moose population according to common practice in Sweden, the hunting was adjusted to keep the population at approximately 65% cows and 35% bulls and to fulfill the condition that 50% of moose felled are calves in all scenarios.

## Results and discussion

We analyzed the effects of different forest management and moose scenarios based on the resulting forest distributions, the moose population, and the numbers of moose felled. Currently (2018) the moose population in the MMA is approximately 2000 moose, ≈ 0.84 moose per km^2^ (Älgförvaltningsplanen ÄFO 8, Länsstyrelsen diarienr. 218-15147-2017). Current forest distribution is shown for pine in Fig. 6.

By varying the target moose population according to the different moose scenarios (Table 1) we find that although the number of moose felled increases with the population size as expected, it reaches a maximum sustainable yield of around 780 felled at a population size around 5000 (2.1 moose / km^2^). The population size can be increased further by reducing hunting (Low-hunting scenario Table 1), but this does not increase the numbers felled (Fig. 7). This is explained not only by reduced hunting but also by a reduction in fecundity (less calves per cow) with increasing population due to competition for food (Fig. 5). The maximum population size possible without hunting is around 10000 (4.2 moose/ km^2^). Although even higher densities have been observed in winter, this moose density is higher than observed mean yearly densities in Scandinavia (Lavsund et al. 2003), which may be due to the lack of hunting, or that we do not yet account for migration or effects of food deficiency on survival.

**Figure 7.**
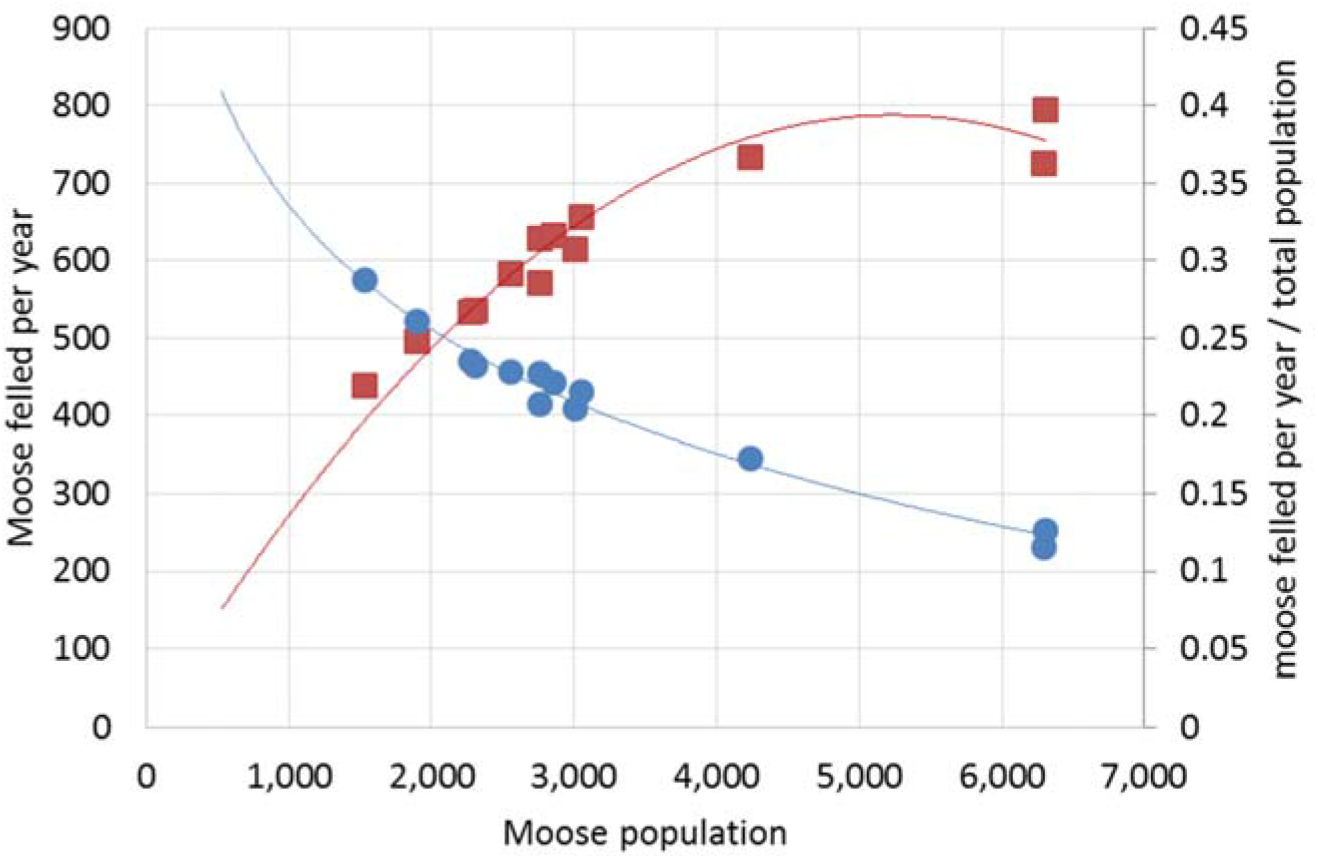
Moose felled (red points and fitted curve) and population productivity (fraction of moose felled per year, blue points and fitted curve) versus total moose population for all different moose management scenarios except no-hunting in the moose management area (Fig. 6, Table 1).

An increasing moose population leads to increased browsing of deciduous trees and pine, which in turn leads to a trade-off between moose population and the productivity of these tree species (Fig. 8). The results also suggest that with a moose population beyond 3000 moose (1.2 moose/ km^2^), additional moose do not reduce the growing stocks of pine further. This result, however, requires further analysis because it may be due to the limited time horizon considered in our scenarios. Because our scenarios run only 20 years the capture effects of moose browsing on young forests but not the subsequent impacts of these damages over the lifetime of the trees. This points to the importance of the time horizon considered and suggest that to understand the full consequences of different scenarios it is necessary to consider longer time horizons.

**Figure 8.**
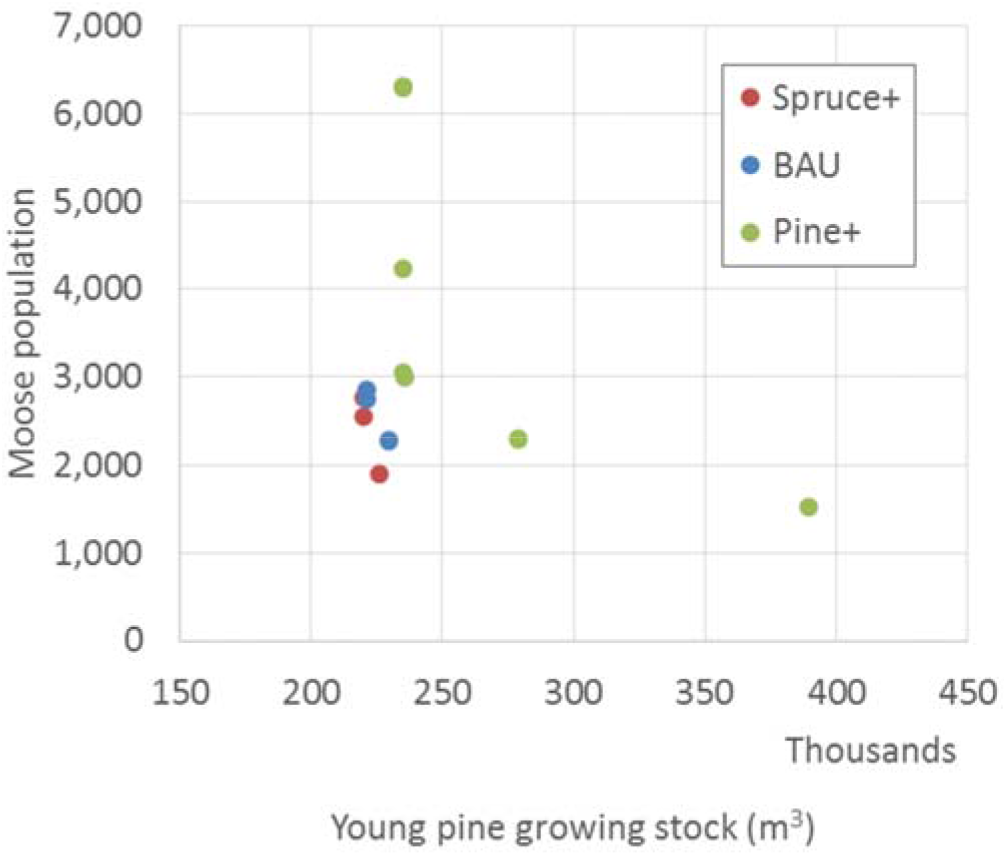
Moose population versus stocks of pine trees with age < 40 years in the moose management area (Fig. 6). The color of points indicates forest management scenario and different points with the same color represent different moose management scenarios (Table 1). The points of the Pine+ scenario suggest a Pareto front on which reduced hunting cannot increase the moose population without reducing pine stocks and increased hunting cannot increase pine stocks without reducing the moose population.

As expected, changing the forest management in terms of planting preferences leads to changes in the forest distribution. A preference for pine (scenario Pine+) where pine is planted on 5% of harvested areas with previously other species leads to small increases in growing stock and spatial distribution of young pine forests (Fig. 9a). Interestingly, the geographical distribution of young pine stocks (age < 40) is shifted over time from west to east. However, for even younger stands up to only 20 years of age, which roughly correspond to age classes browsed by moose (Fig. 3a), the shift is in the opposite direction (Fig. 9b). This means that older stands in the west have been harvested and replaced with young browsable stands and many browsable stands in the east have grown out of the browsable age range. This means that the moose are expected to shift their distribution within the MMA towards more abundant young pine forests in the west during the next 20 years.

**Figure 9.**
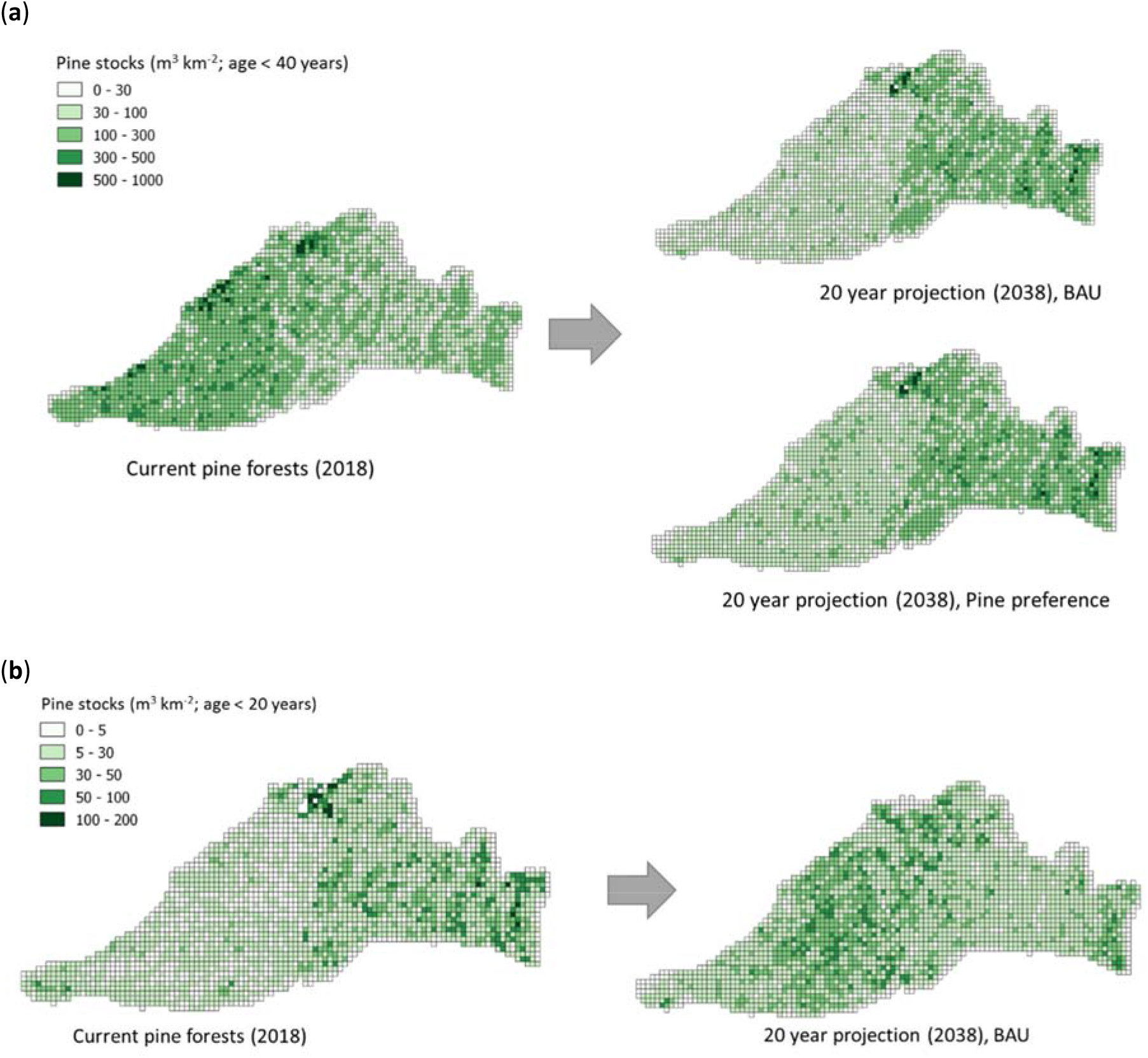
The effect of different forest management scenarios (Table 1) on spatial distribution of young pine forests in the MMA. The moose population was kept at approximately current level (2000 moose, Table 1). (**a**) Pine forests with age < 40 years. (**b**) Pine forests with age <20 years, which correspond approximately to the size range browsed by moose (moose food).

## Conclusions

We conceptualized and developed an integrated model for wildlife and forest management in a group modelling process with modellers (IIASA) and stakeholders (SEPA and NFA). In the process we also identified and addressed knowledge gaps and data needs.

A prototype integrated model of spatially explicit forest and moose dynamics was developed in the project. Although not yet ready and validated for use in real practical applications, the model was shown to be suitable for analyzing the consequences and trade-offs between different goals, such as the maximum sustainable yield of felled moose, and the trade-off between pine forest stocks and the moose population. We also demonstrated how high-resolution spatial forest information can be used for estimating local site index (productivity), perform model calibration, and project shifts in forest distribution over time. The analysis also served to identify limitations and needs for further analysis, such as the importance of the time frame of analysis and the need for characterization of current forest management practices.

Importantly, the project provided a platform in terms of a collaborative network, knowledge, datasets, and key model components, needed for a next stage project to build the comprehensive and refined modeling tools needed for full scale policy support in practice.

## Acknowledgements

Financial support was provided by the Swedish Environmental Protection Agency (SEPA). Maria Hörnell Willebrand and Hördur Haraldsson at SEPA and Christer Kalén at the Swedish Forest Agency (NFA) contributed expertise and information.

